# S-Wipe: stool sample collection for metabolomic gut health tracking

**DOI:** 10.1101/2024.04.12.589313

**Authors:** Alexey V. Melnik, Konstantin Pobozhev, Ali Lotfi, Dana Moradi, Hannah Monahan, Evguenia Kopylova, Alexander A. Aksenov

## Abstract

Microbiome is increasingly recognized as a key factor in health. Intestinal microbiota modulates gut homeostasis via a range of diverse metabolites. Molecules such as short chain fatty acids (SCFAs), the microbial fermentation products of dietary fiber, have been established to be reflective of microbiome and/or dietary shifts and have been linked to multiple gastrointestinal disorders from cancer to colitis, and thus present an excellent diagnostic target. Yet, technical bottlenecks preclude broad translation of such established biomarkers into routine medical practice. In particular, easily accessible, reproducible and robust sampling of stool remains challenging. Here we present Stool Wipe (S-Wipe), an ultra low cost, simplified fecal specimen collection approach designed to overcome key translational barriers without compromising analytical rigor. This sampling approach harnesses lint-free mass spectrometry-compatible cellulose wipes used as a regular toilet paper. The collected stool specimens are then preserved in ethanol solution, do not require refrigeration and can be shipped via regular mail. Using mass spectrometry, we have demonstrated a broad range of captured metabolites, both volatile and non-volatile. The reproducibility and stability of the method was validated for a panel of molecules of particular diagnostic interest, including SCFAs and p-cresol. We demonstrate sensitivity as well as stability and reproducibility of various metabolites collected with S-Wipe. We further demonstrate that S-Wipe is equivalent to the direct stool collection and thus could be used interchangeably and compared to other studies where stool is collected directly. This methodology is ideally suited and is scalable for broad population-based studies, longitudinal tracking such as therapeutic interventions and personalized medicine.

**IMPORTANCE:** Gut microbiome and intestinal metabolome present invaluable diagnostic and therapeutic targets. However, conventional stool testing has several barriers limiting bioassessment from populations. Routine, high temporal resolution monitoring of stool metabolome, including validated biomarkers such as SCFAs, is not implemented due to relatively high cost and inconvenience of sampling, possible need for clinical setting for sample collection, difficulty to collect samples reproducibly, especially due to possible user errors, requirement for freezer storage and maintaining cold chain during shipment. We present a sampling strategy specifically designed to overcome these obstacles. This method can enable capturing accurate molecular snapshots at massive scales, at ultra low cost. The approach collapses complex medical-grade collection into easy self-administration. Individuals can thereby self-monitor therapeutic responses through routine metabolome tracking, including the volatilome, otherwise hindered by infrastructure restrictions. Ultimately, this sampling approach is intended to enable participatory wellness transformation through practical high frequency self-sampling.

---

The gut microbiome plays a central role in human health in profound and wide-ranging ways through nutrient metabolism, immune modulation, and metabolic regulation, including production of bioactive metabolites (1–3). Through production of molecules like short-chain fatty acids (SCFAs), gut bacteria influence processes spanning digestion, immunity, metabolism, and even mental health (4–6). SCFAs can be measured across the gut, with fecal samples being the most accessible (7–10). SCFAs are predominantly produced through fermentation of dietary fibers, subsequently permeate throughout the body, and affect a diverse range of tissues and physiological processes (10–12). The clinical potential of these microbial metabolites as measurable biomarkers continues to be unraveled across investigations - from IBD, all the way to the inflammatory cascade and metabolic syndrome underpinning contemporaneous epidemics of obesity and cardiovascular disease afflicting aging populations (13–17). SCFAs deficiencies have been associated with low-fiber intake diets, altered gut permeability and inflammation, thereby promoting insulin resistance, elevating risk factors, and increasing probability of cardiovascular events (18–21). SCFAs quantification and profiling offers clinical opportunities for non-invasive monitoring, not only for disease diagnostics, but potentially early risk assessment and exposure-response monitoring, as well nutritional or therapeutic gut microbiome modulation.

However, capturing translational insights into these molecular distributions is challenging, as the biological, interpersonal and technical variations tend to obscure molecular trends. Overcoming inherent irreproducibility requires large-scale population measurements where different influencing factors could be assessed. Such large-scale sampling poses practical barriers. Fecal samples rapidly degrade in ambient conditions due to chemical degradation and microbial growth, while reliance on specialized collection and handling infrastructure, cost and complexity of collection, coupled to a wide range of non-standardized sampling protocols poses logistical challenges. Consequently, at the present time, both patients and the general public do not have access to reliable personalized monitoring of gut molecular biomarkers. Last, but not the least, collecting feces is objectionable to many people, limiting their willingness to participate in studies that require fecal testing, especially those that entail frequent longitudinal collection.

Current stool sampling collection methodologies for fecal sampling for microbiome and metabolomics analysis used in clinical practice are predominantly traditional stool collection kits consisting of stool hats, swabs, or small containers (22–28). Samples must be immediately frozen and then transported frozen to a lab to help preserve their integrity. This method can have high variability due to differences in storage and transportation temperature, transit time and exact collection methods between patients. Another common method is fecal occult blood test (FOBT) cards (29–33). In FOBT sampling fecal smears are collected at home on specialized sample cards designed to detect blood. Those cards help stabilize some molecules for transport at room temperature. However, the small sample size and difficulties in using and handling cards limit broader profiling. Specialized solutions such as the Cologuard® can be used for non-invasive tests that analyze stool samples patients collect on a commercial kit with a preservative buffer. This test uses genetic sequencing to detect colorectal precancers but is not optimized for metabolomics (32–34). For the majority of stool-based tests (screening for blood in the stool, stool culturing for bacterial or viral infections diagnostics, ova and parasite exam, fat malabsorption assessment, etc.), stool is collected in research labs or hospitals, where clinicians may collect fecal samples directly, or during procedures like lavage or endoscopic biopsies. Obviously, such invasive techniques limit practical scalability (32).

Overall, sampling methods that are scalable, patient- and consumer-friendly and preserve integrity during non-refrigerated transportation, have remained an unmet need. Most testing today still relies on traditional immediate frozen collection protocols (22, 24), which are logistically impractical for population-level studies. To bridge this gap, we developed the S-Wipe, an ultra low-cost fecal collection method. Cellulose collection paper used in S-Wipe acts as lavatory towels, which makes sampling identical to a normal bathroom routine. Using GC-MS and LC-MS, we demonstrate that S-Wipe captures native SCFAs and other diverse diet-/host-associated metabolites at highly reproducible levels over weeks, without cold storage. The S-Wipe method is designed to enable democratized, scalable gut molecular monitoring to stimulate personalized solutions across global populations.

## RESULTS

The S-Wipe sample collection matrix utilizes cellulose adsorbent paper to passively capture fecal material when used as conventional toilet paper during normal bathroom routines. Approximately 100 mg of stool can be typically collected. The wipe is then placed by the user into a 20 mL wide-mouth S-Wipe funnel or 50 mL centrifuge tube with 5mL of MS-grade 60% ethanol spiked with internal standard, sealed, and then can be both stored and/or shipped at room temperature until extraction and analysis (Figure 1).

**Figure 1.**
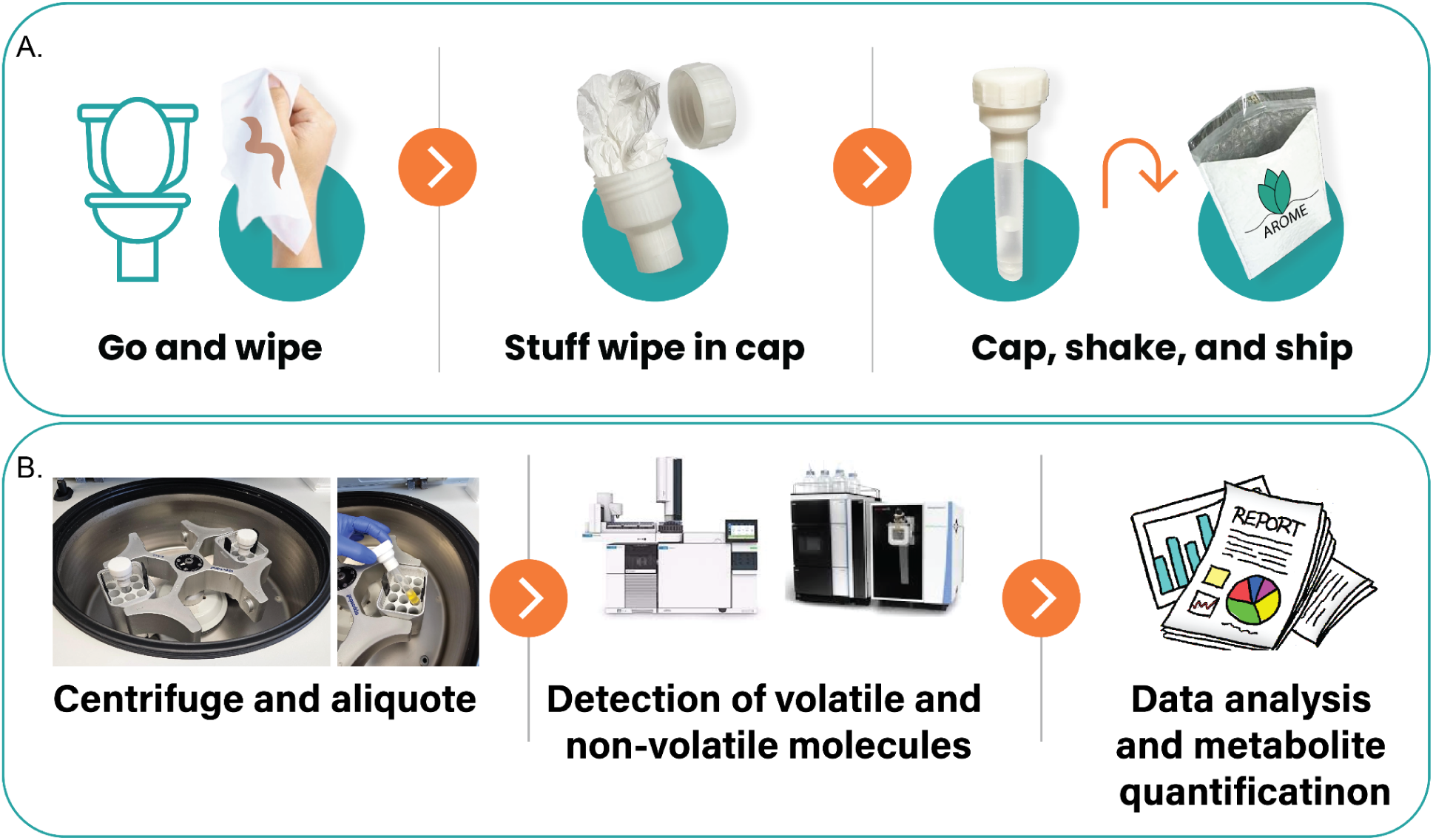
S-Wipe* sampling. a) Sampling and shipping workflow using the kit (Sample collection kit is shown in Supplementary Figure 1). The users are not required to modify their bathroom routine with the exception of placing the user paper into a collection tube rather than flushing/throwing it away. Due to simplicity of collection, every bowel movement could be sampled, if needed, and large longitudinal sampling with each subject being their own control for intervention studies are facilitated. b) Schematic sample preparation and analysis workflow. During shipment, the extraction solvent remains absorbed by the wipe. Upon arrival at the analysis lab, the samples are centrifuged to separate supernatant from the wipe (the cap has a coarse mesh to retain the wipe in the cap). The supernatant is compatible with mass spectrometry analysis, such as GC-MS for SCFA measurement, without any additional sample preparation steps. The absolute measured abundances of biomarkers of interest, as well as longitudinal trends can then be used to inform users. *The patent application is pending (US Application #63/570,322)

The S-Wipe sampling was found to capture the compounds of interest such as SCFA, p-cresol and other molecules that are expected to be detectable in stool, such as indole, bile acids, vitamins, amino acids, lipids etc. (Protocol Optimization study, see Methods)(Supplementary Figure 2 and Supplementary Figure 4). As S-Wipe allows for measuring the stool weight, the abundances of metabolites could be normalized (r(Pearson) = 0.95, p < 0.001, Figure 2A)(Stability study, see Methods). Also, with the use of external calibration, absolute quantities for various molecules could be calculated and used as biomarkers to gain lifestyle or other information, e.g. Supplementary Figure 2A,B. (Interpersonal Variability Study, see Methods) The ratios of metabolites are not dependent on the absolute quantitation, and can also be linked to specific conditions (35–46, 39) (Supplementary Figure 2D).

**Figure 2.**
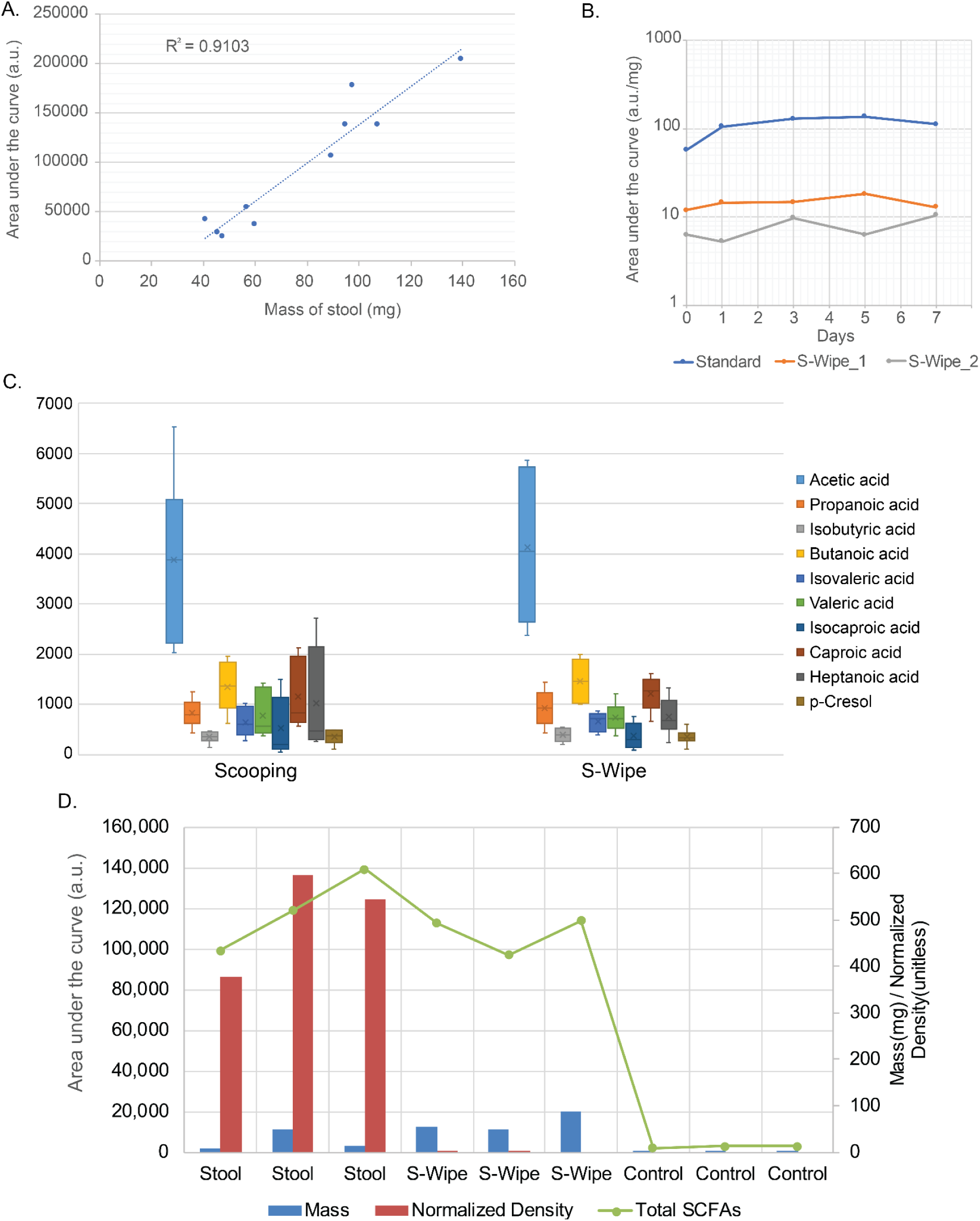
A) Scatter plot of total abundance of C2-C7 SCFAs vs. the measured mass of collected samples for both stool and S-Wipe. A significant positive correlation ((Pearson’s R = 0.95, p < 0.001)) is evident. B) Stability of SCFAs. A 500μl aliquot of the Accustandard FAMQ-004 standard at 10mM concentration was spiked onto S-Wipe. The wipes spiked with standard, and two sampled wipes were then stored for times ranging from one day to one week at room temperature. GC-MS analysis was conducted for each time point to measure abundance of SCFAs. The observed lack of pronounced changes in abundance is suggestive of analyte’s stability within one week time frame. C) Comparison of variability in SCFAs for stool collected with scooping method (conventional sample collection) vs. S-Wipe collection. Sample collected by both methods was extracted and abundances of three SCFAs of primary diagnostic interest (left) and other SCFAs and p-cresol (right) were measured by GC-MS. For all metabolites, the measured abundances generally align, while variability is consistently lower for S-Wipe collection. The variability of acetic acid is greatest due to its highest volatility. D) Plot showing the total abundance of SCFAs, collected using regular stool collection and S-Wipe (green trace). Plot is also showing mass of stool collected in milligrams on the secondary axis (blue bars) as well as Normalized density (ND) of samples as determined by CellsBin proprietary technology (description is given in the Methods section) of real time cell counting (Supplementary video and data). It is evident that the abundance of metabolites is similar between conventional stool collection and S-Wipe, but the cells are generally absent in S-Wipe samples, ostensibly due to their retention on the tissue.

Stability achievable with the S-Wipe was demonstrated for 9 endogenous short chain fatty acid peak areas over the course of 7 days for samples stored at room temperature. The average relative standard deviation error was found to be 23.48% for all time points of 2 replicates (Stability study, see Methods)(Figure 2B). The sensitivity of the approach was on par with conventional stool collection, while sample-to-sample variability is comparable or lower (Figure 2C). Quantitative linearity of short-chain fatty acid detection spanned physiologic concentrations (r2 > 0.99)(Supplementary Figure 3). Absolute concentrations aligned with values reported from conventionally-collected samples, confirming compositional integrity.

In the proposed approach, 60% ethanol solution is used for sample preservation. The ethanol is known to both denature enzymes, preventing enzymatic degradation (47–51), and kill microorganisms preventing bloom (47, 51–53). Focusing on the SCFAs, it is evident that S-Wipe-collected organic acid remained stable for at least several days at ambient temperature. GC-MS indicated negligible changes, with no statistically significant degradation trends over the entire interval (p > 0.24) (Figure 2B).

To more rigorously assess the stability of metabolome, we have further conducted a shipping/storage experiment, where the samples were compared between different handling conditions: refrigerated storage, room temperature storage, short- and long-distance shipping (Figure 3). Correspondingly, the samples collected by S-Wipe were stored at −80C immediately after collection, stored at room temperature for the duration of the study (1 week), shipped, then returned, locally (within the state) and shipped/returned long distance (from the East to West coast of the US). The shipping reflects representative sum total fluctuations in ambient conditions, in particular exposures to high temperatures and humidity. The measured abundances of SCFAs are consistent across all three sample handling scenarios, indicating that their degradation did not occur in a noticeable manner (Figure 3B). Consequently, such a handling strategy is at least appropriate within the outlined conditions for the molecules of interest described here.

**Figure 3.**
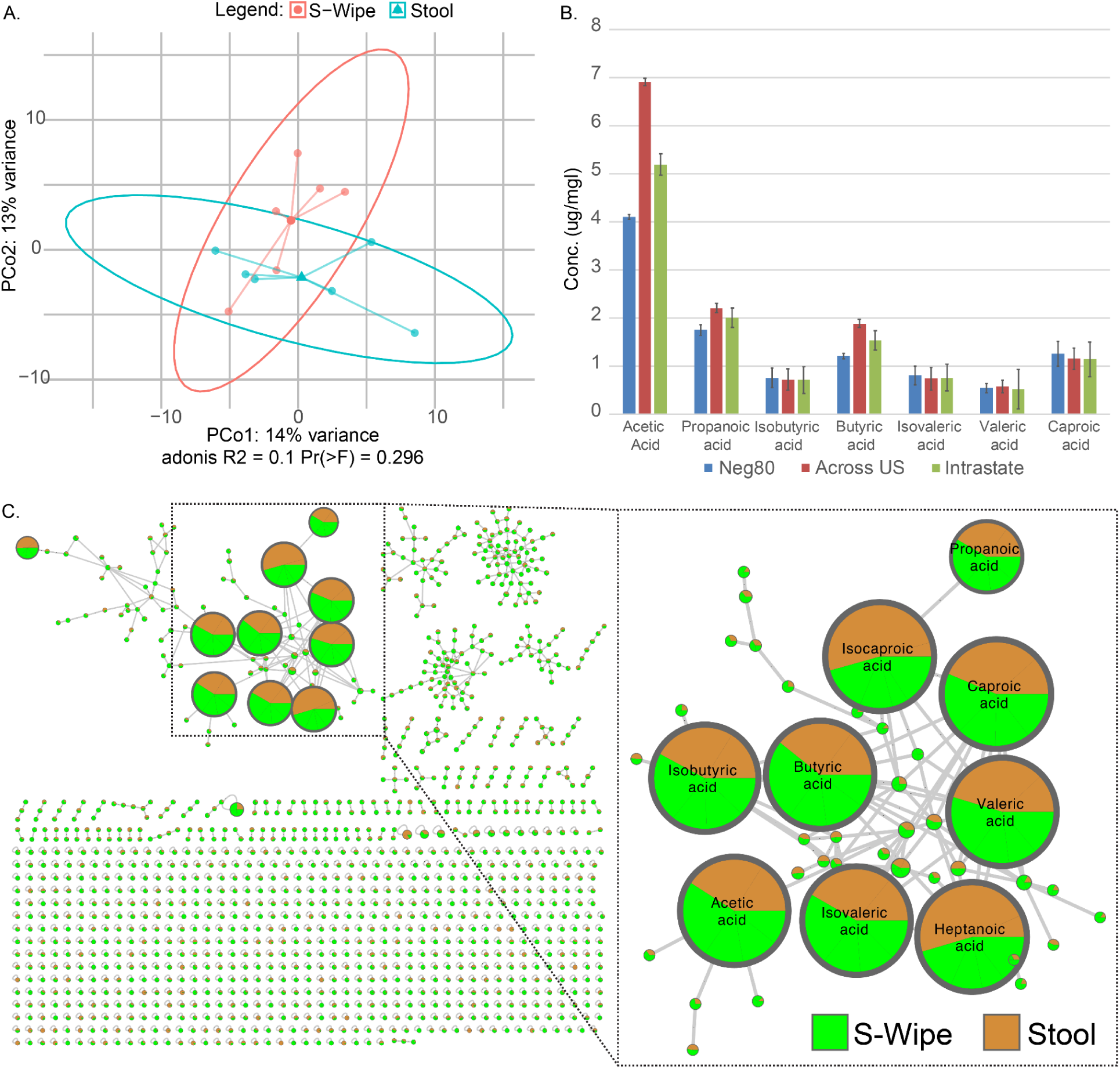
A) Principal coordinate analysis plot of samples from the conventional stool collection and S-Wipe. Several extraction protocols have been tested, as described in the Methods section, to account for possible biases due to the sample preparation. No statistical difference between methodologies is notable. B) Bar plot showing the distribution of short- and medium-chain fatty acids as well as p-cresol and phenol across three different conditions in the shipping study: Neg80 - samples stored at −80C immediately after collection; Intrastate - samples shipped within the state (approximately one week passed in between shipping and receiving samples); Across US - samples shipped from East to West coast of the US (approximately two weeks passed in between shipping and receiving samples). The “Intrastate” and “Across US” samples were stored at 2°C-4°C until analysis. All of the samples were analyzed by GC-MS at the same time. C) Molecular co-network of GC-MS data collected using S-Wipe and regular sample collection approaches; an inset shows clusters of SCFAs. The pie-chart coloring for each molecule’s node corresponds to an averaged abundance of the molecule across all samples collected with the corresponding method; the abundances of various SCFAs are generally comparable and consistent for both methodologies.

The metabolome captured by the S-Wipe was further broadly compared to that of conventional stool collection methods for both volatile and non-volatile molecules. Principal component analysis (PCA) plots revealed no statistically significant differences between the two collection methods (Figure 3A). Further investigation using molecular networking (54, 55) confirmed that the biogenic molecules were consistently captured by S-Wipe in the same way as for the direct stool collection (for example, Figure 3C and Supplementary Figure 4). All examined noted differences between the collection methods were found to correspond to background molecules present in blank samples (Supplementary Figure 5). This corroborates that the S-Wipe methodology collects stool samples without alteration and, with appropriate normalization and blank subtraction, should be directly comparable to any other methodology that does not alter stoll metabolome upon collection.

To confirm the suitability of the S-Wipe method for streamlined metabolomics workflows, we assessed its compatibility with mass spectrometry (MS) analysis without the need for extensive sample manipulation and cleanup. In MS-based metabolomics, various methodologies such as lyophilization/reconstitution, solid-phase extraction, and bi-phase extraction are commonly employed to remove unwanted background components, including proteins, lipids, and cell debris, which can interfere with MS analysis by causing clogging, introducing background noise, and affecting analyte solubility and ionization efficiency (56–58).

Fecal samples naturally contain a large number of host and microbial cells that must be removed before introduction into the MS system (58–60). Unlike direct fecal sample collection methods, the S-Wipe approach is designed to retain the fecal material within the paper matrix while allowing soluble metabolites to partition into the solvent. We hypothesized that this feature of the S-Wipe method would result in a lower cellular content in the collected samples compared to traditional fecal collection techniques. To test this hypothesis, we conducted a comparative cell counting analysis between samples obtained using the S-Wipe approach and those collected through conventional methods. The results confirmed that specimens collected with the S-Wipe approach contained substantially fewer cells (Figure 2D). Correspondingly, by minimizing the presence of cell debris, the S-Wipe method mitigates the risk of instrumentation contamination and circumvents issues such as clogging that may arise from excessive cellular material. The reduction in cellular content thus enables simplified extraction protocol for S-Wipe samples described in the Figure 1B, where a single centrifugation step yields a supernatant that is ready for direct MS analysis.

## DISCUSSION

The motivation behind the present work is to develop an easy-to-use and inexpensive method for stool sampling, focusing on capturing metabolites important for health diagnostics, especially short-chain fatty acids (SCFAs). Such methodology is essential for large-scale projects that explore the metabolome in conjunction with human gut health. Combination of food molecules, exposures, host’s endogenous metabolites, and their various transformations, including microbially-contributed metabolites, make gut some of the most molecularly complex environments. To capture the intricacies of native biological complexity and account for the variability in the general population, studies must be conducted on a fundamentally different scale, involving large, diverse cohorts. In practical terms, this necessitates the collection and analysis of an immense number of samples, ranging from thousands to ultimately millions or even billions.

Revolutionary advances in sequencing two decades ago have enabled previously unimaginable magnitudes in studies such as the Human Microbiome Project (61–68) and the Earth Microbiome Project (EMP) (69–71). The latter project intends to capture the full microbial diversity of the entire planet. Studies of this magnitude must be community-driven, as no single lab would be able to conduct such research by itself. Harmonized, large-scale sampling is central to making such efforts feasible, as it allows multiple laboratories to collect samples independently and then conduct joint co-analysis of collected data (61–63, 71).

Unlike in DNA sequencing, metabolomics sampling approaches are generally neither standardized nor optimized for broad-range molecular explorations. This is due to the inherent difficulty to standardize sample collection in MS, and, concurrently, the current lack of truly large-scale efforts of population-wide studies similar to those done by 16S or metagenomics sequencing (64)(72). This creates a negative feedback loop, where the lack of established standardized sampling methodologies complicates large-scale metabolomics studies, and conversely, the lack of large-scale studies comparable in ambition to EMP does not provide the impetus to develop sample methodologies appropriate for population-wide scale studies.

A major factor in lack of sampling standardization in metabolomics is due to the fact that, unlike in sequencing, where DNA properties are relatively consistent across samples, molecular properties vary widely, as molecules differ in size, polarity, volatility, solubility, stability, etc. Standardizing any one methodology inevitably creates the possibility of bias, where a subset of the metabolome is efficiently detected, but some molecules are either not collected, or lost during the sample handling, or would not be detectable with the sample processing (e.g., due to poor solubility). A methodology choice is often dictated by the research question at hand and the molecules of interest in each individual study. In particular, the choice of solvent is crucial, as it selectively dissolves certain molecules and directly affects the detectable metabolome.

Aqueous solvents are more suitable for detecting polar molecules, while organic solvents are better for detecting less polar compounds. Considering these limitations, a “universal methodology” in metabolomics would be different from that in DNA sequencing. Such standardized sampling methodology cannot be absolutely uniform, but instead it needs to: 1) collect native samples to preclude bias due to the sampling itself; 2) avoid or minimize method-specific distortion due to sample handling, preparation etc. of detectable molecular distributions; and 3) be amenable to a variety of analysis methodologies. The choice of methodology will then dictate the downstream sample manipulation. In the context of metabolomics, the main methodologies are predominantly GC-MS (73–75) and LC-MS (76–80), and less often, NMR (81–83). Additionally, the sampling methodology should be inexpensive, simple to use to minimize user error, widely accessible (including minimizing or, preferably, eschewing the use of critical materials, particularly metals (56, 84–89), and scalable).

The S-Wipe methodology was developed with the above considerations in mind. This sampling manifold is cheap (<$1 per kit), easy to manufacture, and uses broadly available components and materials with minimal design complexity. More importantly, the approach is intended to minimize user burden, enabling individuals to adopt it without disrupting their established routines. Using the toilet wipe allows collecting neat fecal samples without alteration, as evidenced by the results presented above (Figure 3A and Supplementary Figure 4). Since the samples collected by S-Wipe are directly comparable to neat stool samples, data from any study utilizing S-Wipe, in principle, should be directly comparable to studies where neat stool is collected by any other means, enabling continuation of existing studies, building biobanks, routine sampling, cross-laboratory data comparison, etc., provided standardization of downstream analysis protocols.

In this study, 60% ethanol was chosen as a “middle ground” solvent. It is suitable for sample preservation during shipping and for capturing and detecting many important metabolites, particularly short-chain fatty acids (SCFAs). Ethanol is also convenient due to its relatively low toxicity, cost-effectiveness, and suitability for a broad range of mid- to low-polarity molecules. We suggest using 60% ethanol for gut metabolome studies in particular, as we found convincing evidence that the degradation of the SCFAs is negligible, even when stored at room temperature for multiple days (Figure 2B). While SCFAs demonstrated stability under the studied conditions, it is important to note that other, less stable molecules may still be susceptible to degradation. Therefore, when considering other molecules for diagnostic purposes, it is crucial to establish their stability, if not already known, in order to ensure the reliability and accuracy of the results obtained using the S-Wipe methodology.

Ethanol denatures enzymes and kills off microflora, thus minimizing the biochemical alteration of molecules in the sample and preventing microbial blooms. However, ethanol may not be the best choice for all applications. For instance, other solvents may need to be used in addition to or instead of ethanol, depending on the specific application. Use of certain solvents is necessary for optimizing the detection of particular molecules or molecular classes. The S-Wipe sample preparation workflow could be adopted accordingly. By lyophilizing the sample and reconstituting it in a different solvent, a broad range of molecules present in the sample can be detected (this is only applicable for non-volatile molecules, as lyophilization step will remove or distort abundances for volatile and semi-volatile molecules). This allows for the optimization of the solvent for compatibility with the analytical methodology (e.g., RP vs. HILIC for LC-MS) based on the specific compounds of interest. Since the use of different downstream sample handling will result in the detection of different molecular distributions, it is essential to standardize the use of the same solvent throughout each study or set of studies to ensure consistency and reproducibility across.

The key advantage of the S-Wipe method is the minimal complexity of sample acquisition, so any able person can use it without requiring specialized facilities or equipment. The amount of material can vary with each use, but the collected sample weight can be directly measured. Both the weight of the collection tube with solvents and the weight of the wipe are standardized, allowing for the determination of the collected sample weight by subtracting these known values from the weight of the used kit. Fecal metabolite concentrations often demonstrate poor direct correlations to total stool mass due to considerable individual variations in water content and dietary fiber, as the water and non-fermented fibrous fraction expands stool weight without affecting bioactive compounds. With the use of S-Wipe, we observed a strong correlation between the total amount of SCFAs and biomass, indicating the utility of normalization by mass (Figure 2A).

Finally, we note the minimal presence of cells in the collected samples. This further makes the methodology conducive to high-throughput metabolomic profiling by alleviating the need for extensive sample cleanup, rendering this novel collection strategy well-suited for large-scale studies.

Taken together, the results demonstrate that the S-Wipe method collects and preserves stool metabolites that could be used in health assessments. The methodology itself is a very simple to use, cost-effective, and versatile approach to stool sampling for metabolomics analysis. By enabling the collection of native samples without bias, minimizing method-specific distortions, and allowing for adaptability to various analytical methodologies, this method has the potential to facilitate large-scale, population-wide gut metabolome studies. The standardization of sample handling and the use of appropriate solvents, such as 60% ethanol, would ensure consistency and reproducibility across studies. With its ease of use and minimal sample cleanup requirements, the S-Wipe method is well-positioned to support the growing need for large-scale metabolomics investigations in the context of human gut health and beyond.

## Conclusion

This study describes a fecal sampling routine designed for high-frequency metabolomics sampling. The approach condenses into the ubiquitous toilet wipe use without compromising analytical sensitivity or quantitative molecular preservation. We have shown the intrinsic capacity to stably sorb, store, and release key gut metabolites such as SCFAs and Bile acids using an inexpensive, immediately accessible paper matrix suitable for self-administration. By simplifying the process, this approach is designed to translate monitoring of the gut microbiome on the molecular level from an academic curiosity into a practical tool for personalized nutrition and disease management. Similar to how affordable genetic testing has allowed proactive mitigation of hereditary risks, routine measurement of molecular biomarkers such as SCFAs has a potential to shift gastrointestinal disease management from reactive to predictive by providing easy access to gut metabolome monitoring.

## Materials and Methods

### Protocol Optimization Study

A healthy volunteer donor has provided all stool samples. The protocol described in the “Sample Extraction” section was run alongside a long sonication and incubation extraction protocol. The long protocol included a 10 minute sonication on ice and six hours on a vibration table at 2 - 4°C for homogenization. Samples included S-Wipe kits with no wipe, S-Wipe kits complete, and S-Wipe kits with no stool. Samples were generated in triplicates. After sample extraction, samples were analyzed by GC-MS and LC-MS/MS.

### Stability Study

A healthy volunteer donor has provided all stool samples. Samples collected by S-Wipe kits and controls were incubated at 25°C for 0-, 3-, 5-, and 7- days. Negative controls contained no stool and positive controls contained no stool and Accustandard FAMQ-004 at 1mM. All samples were collected in triplicates. Sampled kits were stored at −80°C until analysis and all samples were extracted together. After sample extraction, samples were analyzed by GC-MS and LC-MS/MS.

### Shipping Study

A healthy volunteer donor has provided all stool samples. Samples collected by S-Wipe kits and controls were sent and returned via United States Postal Service to a short-distance (36-miles) and long-distance (2,950-miles) address. Negative control contained no stool. All samples were in triplicates. Upon reception in the mail, the S-Wipe kits were stored at 2 - 4°C until analysis. All samples were extracted and analyzed by GC-MS and LC-MS/MS at the same time.

### Interpersonal Variability Study

10 healthy volunteers were provided with S-Wipe collection kits and US mail return labels to return samples to the lab. No personal or any other information was collected throughout the study.

### Sample Extraction

Completed S-Wipe kits kept at −80°C were thawed on ice prior to extraction. Pre-sample and post-sample kit mass was recorded. The completed kits were first placed in an iced ultrasonic bath for 10 minutes and then on a vibration table for 10 minutes at room temperature for homogenization. 1000μL aliquot was taken and transferred to 1.5 mL microcentrifuge tubes. The samples were centrifuged for 5 minutes at 14,000 revolutions per minute. After centrifugation, aliquots of 100 uL were taken from each sample and transferred to vials with conical inserts, and analyzed by GC-MS and/or LC-MS/MS. Samples were directly loaded onto GC-MS. For LC-MS/MS samples were diluted 4-fold with pure cold methanol to precipitate protein. Samples then were filtered through Phenomenex Phree plates with the application of 4 psi negative pressure. 200 uL of collected filtrates were dried under vacuum and reconstituted with 100uL of C18 resuspension buffer (5% Acetonitrile (Sigma, USA) in LCMS grade water with added internal standards: Sulfachloropyridazine (TCI, USA) and Sulfamethazine (Sigma, USA)).

### GC-MS data acquisition and processing

1 μL aliquot of the S-Wipe kit supernatant was directly injected into an Agilent 6890 GC interfaced to a mass spectrometer Hewlett Packard MSD 5973 for electron ionization GC-MS. The GC utilizes a 30m ZB-FFAP column (0.25 mm i.d., 0.25 μm film thickness) for metabolite separation with 1.2mL/min constant He flow. The oven temperature program initiates at 50°C rising to 240°C at 10°C/min. Lint particulate traces from wipe material were found to contribute no observable analytical interference. The data were then deconvoluted with the MSHub algorithm (54). The experimental spectra were searched against the NIST 2023 library with ≥80% spectral match defining putative identifications, and the retention times falling within <0.01 min of the corresponding reference standards. Targeted analysis of SCFAs and p-cresol was performed by using Agilent Masshunter software.

### LC-MS/MS data acquisition and processing

The samples were injected and chromatographically separated using a Vanquish UPLC (Thermo Fisher Scientific, Waltham, MA), on a 100 mm × 2.1 mm Kinetex 1.7 μM, C18, 100 Å chromatography column (Phenomenex, Torrance, CA), 40 °C column temperature, 0.4 mL/min flow rate, mobile phase A 99.9% water (J.T. Baker, LC–MS grade), 0.1% formic acid (Thermo Fisher Scientific, Optima LC/MS), mobile phase B 99.9% acetonitrile (J.T. Baker, LC–MS grade), 0.1% formic acid (Fisher Scientific, Optima LC–MS), with a the following gradient: 0–1 min 5% B, 1–8 min 100% B, 8–10.9 min 100% B, 10.9–11 min 5% B, 11–12 min 5% B.

MS analysis was performed on Orbitrap Exploris 240(Thermo Fisher Scientific, Waltham, MA) mass spectrometer equipped with HESI-II probe sources. The following probe settings were used for both MS for flow aspiration and ionization: spray voltage of 3500 V, sheath gas (N_2_) pressure of 35 psi, auxiliary gas pressure (N_2_) of 10 psi, ion source temperature of 350 °C, S-lens RF level of 50 Hz and auxiliary gas heater temperature at 400 °C.

Spectra were acquired in positive ion mode over a mass range of 100–1500 *m*/*z*. An external calibration with Pierce LTQ Velos ESI positive ion calibration solution (Thermo Fisher Scientific, Waltham, MA) was performed prior to data acquisition with ppm error of less than 1. Data were recorded with data-dependent MS/MS acquisition mode. Full scan at MS1 level was performed with 30K resolution. MS2 scans were performed at 11250 resolution with max IT time of 60 ms in profile mode. MS/MS precursor selection windows were set to *m*/*z* 2.0 with *m*/*z* 0.5 offset. MS/MS active exclusion parameter was set to 5.0 s. Raw MS/MS data were converted to mzML files using MSConvert.

LC-MS raw data files were converted to mzML format using msConvert (ProteoWizard). MS1 features were selected for all LC-MS data sets collected using the open-source software MZmine 3 (90) with the following parameters: mass detection noise level was 10,000 counts, chromatograms were built over a 0.01-min minimum time span, with 5,000-count minimum peak height and 5-ppm mass tolerance, features were deisotoped and aligned with 10-ppm tolerance and 0.1-min retention time tolerance, and aligned features were filtered based on a minimum 3-peak presence in samples and based on containing at least 2 isotopes. Subsequent blank filtering, total ion current, and internal standard normalization were performed for representation of relative abundance of molecular features and for principal-coordinate analysis (PCoA).

### Cell counting assay

Blind samples were analyzed using CellsBin’s VEGA platform at an external lab. The normalized density of each sample was calculated by imaging 20 µl through CellsBin’s microfluidic optical device. The company’s machine learning algorithm detected and classified particles, estimating the relative size and normalized density of each sample.

## Supporting information

Supplementary_Video_sample-1

Supplementary_Video_sample-2

Supplementary_Video_sample-3

Supplementary_Video_sample-10

Supplementary_Video_sample-11

Supplementary_Video_sample-12

Normalized Density

## Data availability

All data generated in this study are publicly available. The raw data are available on MassIVE Repository (massive.ucsd.edu) under the following dataset accession numbers <u>MSV000094530</u> (GC-MS) and <u>MSV000094529</u>(LC-MS)

## AUTHOR CONTRIBUTIONS

AM and AA formulated the study. AM, KP, HM prepared the samples. AM devised the sampling manifold. AM, KP performed the GC-MS and LC-MS analysis. EK, AM conducted data processing and statistical analysis. AA, AM, HM coordinated the study. AA, AL, DM, AM wrote the manuscript. All authors read and approved the manuscript.

## Competing interests

AA and AM are founders of Arome Science Inc. EK is a founder and director of Clarity Genomics. The authors declare that they have no other competing interests.

## Acknowledgments

The authors appreciate the help and materials in the form of automation support provided by TomTec Inc. (www.tomtec.com). Additionally, the authors thank CellsBin Inc. (www.cellsbin.com) for the analysis of samples for microscopy measurements (CellsBin, based in New Haven, CT, is a tech-bio company. Its VEGA platform is utilized in precision oncology and other fields, including aerosol detection for the prestigious IARPA’s Picard Project (https://www.iarpa.gov/research-programs/picard)). Arome Science Inc. provided funding for the study.

## Notes

### Summary of Updates

Some video added to supplementary, which show Normalized density (ND) of samples as determined by CellsBin proprietary technology of real time cell counting.

## References

1. Duvallet C, Gibbons SM, Gurry T, Irizarry RA, Alm EJ. 2017. Meta-analysis of gut microbiome studies identifies disease-specific and shared responses. Nat Commun 8:1–10.

2. 2018. The Intestinal Microbiota in Colorectal Cancer. Cancer Cell 33:954–964.

3. Shreiner AB, Kao JY, Young VB. 2015. The gut microbiome in health and in disease. Curr Opin Gastroenterol 31:69.

4. Clemente JC, Pehrsson EC, Blaser MJ, Sandhu K, Gao Z, Wang B, Magris M, Hidalgo G, Contreras M, Noya-Alarcón Ó, Lander O, McDonald J, Cox M, Walter J, Oh PL, Ruiz JF, Rodriguez S, Shen N, Song SJ, Metcalf J, Knight R, Dantas G, Dominguez-Bello MG. 2015. The microbiome of uncontacted Amerindians. Sci Adv 1.

5. Dalile B, Van Oudenhove L, Vervliet B, Verbeke K. 2019. The role of short-chain fatty acids in microbiota–gut–brain communication. Nat Rev Gastroenterol Hepatol 16:461–478.

6. Ghosh TS, Rampelli S, Jeffery IB, Santoro A, Neto M, Capri M, Giampieri E, Jennings A, Candela M, Turroni S, Zoetendal EG, Hermes GDA, Elodie C, Meunier N, Brugere CM, Pujos-Guillot E, Berendsen AM, De Groot LCPGM, Feskins EJM, Kaluza J, Pietruszka B, Bielak MJ, Comte B, Maijo-Ferre M, Nicoletti C, De Vos WM, Fairweather-Tait S, Cassidy A, Brigidi P, Franceschi C, O’Toole PW. 2020. Mediterranean diet intervention alters the gut microbiome in older people reducing frailty and improving health status: the NU-AGE 1-year dietary intervention across five European countries. Gut 69:1218–1228.

7. 2021. Gut-microbiota-targeted diets modulate human immune status. Cell 184:4137–4153.e14.

8. Hale VL, Chen J, Johnson S, Harrington SC, Yab TC, Smyrk TC, Nelson H, Boardman LA, Druliner BR, Levin TR, Rex DK, Ahnen DJ, Lance P, Ahlquist DA, Chia N. 2017. Shifts in the Fecal Microbiota Associated with Adenomatous Polyps. Cancer Epidemiol Biomarkers Prev 26:85–94.

9. Gou W, Miao Z, Deng K, Zheng J-S. 2023. Nutri-microbiome epidemiology, an emerging field to disentangle the interplay between nutrition and microbiome for human health. Protein Cell 14:787–806.

10. He J, Zhang P, Shen L, Niu L, Tan Y, Chen L, Zhao Y, Bai L, Hao X, Li X, Zhang S, Zhu L. 2020. Short-Chain Fatty Acids and Their Association with Signalling Pathways in Inflammation, Glucose and Lipid Metabolism. Int J Mol Sci 21.

11. Chambers ES, Viardot A, Psichas A, Morrison DJ, Murphy KG, Zac-Varghese SEK, MacDougall K, Preston T, Tedford C, Finlayson GS, Blundell JE, Bell JD, Louise Thomas E, Mt-Isa S, Ashby D, Gibson GR, Kolida S, Dhillo WS, Bloom SR, Morley W, Clegg S, Frost G. 2015. Effects of targeted delivery of propionate to the human colon on appetite regulation, body weight maintenance and adiposity in overweight adults. Gut 64:1744–1754.

12. Fu J, Zheng Y, Gao Y, Xu W. 2022. Dietary Fiber Intake and Gut Microbiota in Human Health. Microorganisms 10.

13. Raman M. 2019. Nutrition in Inflammatory Bowel Disease (IBD).

14. Kochhar S, Martin F-P. 2014. Metabonomics and Gut Microbiota in Nutrition and Disease. Springer.

15. Burke A, Lichtenstein GR, Rombeau JL. 1998. Nutritional and metabolic advances in inflammatory bowel disease. Curr Opin Clin Nutr Metab Care 1:387.

16. Sezikli M, Akkan Cetinkaya Z, Güzelbulut F, Özgil Yetkin D. 2012. Metabolic syndrome in inflammatory bowel disease: A real relationship or just a coincidence. Euroasian J Hepatogastroenterol 2:79–83.

17. Bilski J, Mazur-Bialy A, Wojcik D, Surmiak M, Magierowski M, Sliwowski Z, Pajdo R, Kwiecien S, Danielak A, Ptak-Belowska A, Brzozowski T. 2019. Role of Obesity, Mesenteric Adipose Tissue, and Adipokines in Inflammatory Bowel Diseases. Biomolecules 9.

18. Candela M, Biagi E, Soverini M, Consolandi C, Quercia S, Severgnini M, Peano C, Turroni S, Rampelli S, Pozzilli P, Pianesi M, Fallucca F, Brigidi P. 2016. Modulation of gut microbiota dysbioses in type 2 diabetic patients by macrobiotic Ma-Pi 2 diet. Br J Nutr 116:80–93.

19. Walker AW, Ince J, Duncan SH, Webster LM, Holtrop G, Ze X, Brown D, Stares MD, Scott P, Bergerat A, Louis P, McIntosh F, Johnstone AM, Lobley GE, Parkhill J, Flint HJ. 2011. Dominant and diet-responsive groups of bacteria within the human colonic microbiota. ISME J 5:220–230.

20. Gomez-Arango LF, Barrett HL, Wilkinson SA, Callaway LK, McIntyre HD, Morrison M, Dekker Nitert M. 2018. Low dietary fiber intake increases Collinsella abundance in the gut microbiota of overweight and obese pregnant women. Gut Microbes 9:189–201.

21. Zhang X, Shen D, Fang Z, Jie Z, Qiu X, Zhang C, Chen Y, Ji L. 2013. Human gut microbiota changes reveal the progression of glucose intolerance. PLoS One 8:e71108.

22. Zhou Z, Zheng Z, Xiong X, Chen X, Peng J, Yao H, Pu J, Chen Q, Zheng M. 2021. Gut Microbiota Composition and Fecal Metabolic Profiling in Patients With Diabetic Retinopathy. Front Cell Dev Biol 9:732204.

23. Guan H, Pu Y, Liu C, Lou T, Tan S, Kong M, Sun Z, Mei Z, Qi Q, Quan Z, Zhao G, Zheng Y. 2021. Comparison of Fecal Collection Methods on Variation in Gut Metagenomics and Untargeted Metabolomics. mSphere 6:e0063621.

24. Meade S, Liu Chen Kiow J, Massaro C, Kaur G, Squirell E, Bressler B, Lunken G. 2023. Gut microbiome-associated predictors as biomarkers of response to advanced therapies in inflammatory bowel disease: a systematic review. Gut Microbes 15:2287073.

25. Li M, Yang L, Mu C, Sun Y, Gu Y, Chen D, Liu T, Cao H. 2022. Gut microbial metabolome in inflammatory bowel disease: From association to therapeutic perspectives. Comput Struct Biotechnol J 20:2402–2414.

26. Metwaly A, Dunkel A, Waldschmitt N, Raj ACD, Lagkouvardos I, Corraliza AM, Mayorgas A, Martinez-Medina M, Reiter S, Schloter M, Hofmann T, Allez M, Panes J, Salas A, Haller D. 2020. Integrated microbiota and metabolite profiles link Crohn’s disease to sulfur metabolism. Nat Commun 11:4322.

27. Fecal Collection Kits and Devices. ZYMO RESEARCH. https://www.zymoresearch.com/collections/fecal-collection. Retrieved 5 April 2024.

28. EasySampler® Stool Collection Kit. https://www.alpco.com/easysamplerr-stool-collection-kit.html. Retrieved 5 April 2024.

29. Kaur K, Zubair M, Adamski JJ. 2023. Fecal Occult Blood TestStatPearls [Internet]. StatPearls Publishing.

30. Fecal Occult Blood Test (FOBT). https://medlineplus.gov/lab-tests/fecal-occult-blood-test-fobt/. Retrieved 5 April 2024.

31. 2020. Hemoccult. HemoCue America. https://www.hemocue.us/hemoccult/. Retrieved 5 April 2024.

32. Mo S, Wang H, Han L, Xiang W, Dai W, Zhao P, Pei F, Su Z, Ma C, Li Q, Wang Z, Cai S, Wang H, Liu R, Cai G. 2021. Fecal Multidimensional Assay for Non-Invasive Detection of Colorectal Cancer: Fecal Immunochemical Test, Stool DNA Mutation, Methylation, and Intestinal Bacteria Analysis. Front Oncol 11:643136.

33. Zeng T, Huang Z, Yu X, Zheng L, Liu T, Tian B, Xiao S, Huang J. 2023. Combining methylated SDC2 test in stool DNA, fecal immunochemical test, and tumor markers improves early detection of colorectal neoplasms. Front Oncol 13:1166796.

34. Scholefield JH, Moss SM, Mangham CM, Whynes DK, Hardcastle JD. 2012. Nottingham trial of faecal occult blood testing for colorectal cancer: a 20-year follow-up. Gut 61:1036–1040.

35. Martin-Gallausiaux C, Marinelli L, Blottière HM, Larraufie P, Lapaque N. 2021. SCFA: mechanisms and functional importance in the gut. Proc Nutr Soc 80:37–49.

36. Otten BMJ, Sthijns MMJPE, Troost FJ. 2023. A Combination of Acetate, Propionate, and Butyrate Increases Glucose Uptake in C2C12 Myotubes. Nutrients 15.

37. Liu H, Wang J, He T, Becker S, Zhang G, Li D, Ma X. 2018. Butyrate: A Double-Edged Sword for Health? Adv Nutr 9:21–29.

38. Nogal A, Valdes AM, Menni C. 2021. The role of short-chain fatty acids in the interplay between gut microbiota and diet in cardio-metabolic health. Gut Microbes 13:1–24.

39. Hiseni P, Snipen L, Wilson RC, Furu K, Hegge FT, Rudi K. 2023. Prediction of high fecal propionate-to-butyrate ratios using 16S rRNA-based detection of bacterial groups with liquid array diagnostics. Biotechniques 74:9–21.

40. Cong J, Zhou P, Zhang R. 2022. Intestinal Microbiota-Derived Short Chain Fatty Acids in Host Health and Disease. Nutrients 14.

41. Parada Venegas D, De la Fuente MK, Landskron G, González MJ, Quera R, Dijkstra G, Harmsen HJM, Faber KN, Hermoso MA. 2019. Corrigendum: Short Chain Fatty Acids (SCFAs)-Mediated Gut Epithelial and Immune Regulation and Its Relevance for Inflammatory Bowel Diseases. Front Immunol 10:1486.

42. Hoch SE, Kirov II, Tal A. 2017. When are metabolic ratios superior to absolute quantification? A statistical analysis. NMR Biomed 30.

43. Saccenti E, Hoefsloot HCJ, Smilde AK, Westerhuis JA, Hendriks MMWB. 2014. Reflections on univariate and multivariate analysis of metabolomics data. Metabolomics 10:361–374.

44. Smith LM, Maher AD, Want EJ, Elliott P, Stamler J, Hawkes GE, Holmes E, Lindon JC, Nicholson JK. 2009. Large-scale human metabolic phenotyping and molecular epidemiological studies via 1H NMR spectroscopy of urine: investigation of borate preservation. Anal Chem 81:4847–4856.

45. Thévenot EA, Roux A, Xu Y, Ezan E, Junot C. 2015. Analysis of the Human Adult Urinary Metabolome Variations with Age, Body Mass Index, and Gender by Implementing a Comprehensive Workflow for Univariate and OPLS Statistical Analyses. J Proteome Res 14:3322–3335.

46. Li BSY, Wang H, Gonen O. 2003. Metabolite ratios to assumed stable creatine level may confound the quantification of proton brain MR spectroscopy. Magn Reson Imaging 21:923–928.

47. Hass JK, Henriquez MC, Churcher J, Hamou H, Morales SR, Melin AD. 2024. Assessing morphological preservation of gastrointestinal parasites from fecal samples of wild capuchin monkeys (Cebus imitator) stored in ethanol versus formalin. Sci Rep 14:3623.

48. Li X-M, Shi X, Yao Y, Shen Y-C, Wu X-L, Cai T, Liang L-X, Wang F. 2023. Effects of Stool Sample Preservation Methods on Gut Microbiota Biodiversity: New Original Data and Systematic Review with Meta-Analysis. Microbiol Spectr 11:e0429722.

49. Marotz C, Cavagnero KJ, Song SJ, McDonald D, Wandro S, Humphrey G, Bryant M, Ackermann G, Diaz E, Knight R. 2021. Evaluation of the Effect of Storage Methods on Fecal, Saliva, and Skin Microbiome Composition. mSystems 6.

50. Smith L, Villaret-Cazadamont J, Claus SP, Canlet C, Guillou H, Cabaton NJ, Ellero-Simatos S. 2020. Important Considerations for Sample Collection in Metabolomics Studies with a Special Focus on Applications to Liver Functions. Metabolites 10.

51. Rius-Sansalvador B, Bars-Cortina D, Khannous-Lleiffe O, Serrano AG-, Guinó E, Saus E, Gabaldón T, Moreno V, Obón-Santacana M. 2023. Stability of oral and fecal microbiome at room temperature: Impact on diversity. bioRxiv.

52. Marquina D, Buczek M, Ronquist F, Łukasik P. 2021. The effect of ethanol concentration on the morphological and molecular preservation of insects for biodiversity studies. PeerJ 9:e10799.

53. Nosková E, Modrý D, Baláž V, Červená B, Jirků-Pomajbíková K, Zechmeisterová K, Leowski C, Petrželková KJ, Pšenková I, Vodička R, Kessler SE, Ngoubangoye B, Setchell JM, Pafčo B. 2023. Identification of potentially zoonotic parasites in captive orangutans and semi-captive mandrills: Phylogeny and morphological comparison. Am J Primatol 85:e23475.

54. Aksenov AA, Laponogov I, Zhang Z, Doran SLF, Belluomo I, Veselkov D, Bittremieux W, Nothias LF, Nothias-Esposito M, Maloney KN, Misra BB, Melnik AV, Smirnov A, Du X, Jones KL 2nd, Dorrestein K, Panitchpakdi M, Ernst M, van der Hooft JJJ, Gonzalez M, Carazzone C, Amézquita A, Callewaert C, Morton JT, Quinn RA, Bouslimani A, Orio AA, Petras D, Smania AM, Couvillion SP, Burnet MC, Nicora CD, Zink E, Metz TO, Artaev V, Humston-Fulmer E, Gregor R, Meijler MM, Mizrahi I, Eyal S, Anderson B, Dutton R, Lugan R, Boulch PL, Guitton Y, Prevost S, Poirier A, Dervilly G, Le Bizec B, Fait A, Persi NS, Song C, Gashu K, Coras R, Guma M, Manasson J, Scher JU, Barupal DK, Alseekh S, Fernie AR, Mirnezami R, Vasiliou V, Schmid R, Borisov RS, Kulikova LN, Knight R, Wang M, Hanna GB, Dorrestein PC, Veselkov K. 2021. Auto-deconvolution and molecular networking of gas chromatography-mass spectrometry data. Nat Biotechnol 39:169–173.

55. Nothias L-F, Petras D, Schmid R, Dührkop K, Rainer J, Sarvepalli A, Protsyuk I, Ernst M, Tsugawa H, Fleischauer M, Aicheler F, Aksenov AA, Alka O, Allard P-M, Barsch A, Cachet X, Caraballo-Rodriguez AM, Da Silva RR, Dang T, Garg N, Gauglitz JM, Gurevich A, Isaac G, Jarmusch AK, Kameník Z, Kang KB, Kessler N, Koester I, Korf A, Le Gouellec A, Ludwig M, Martin H C, McCall L-I, McSayles J, Meyer SW, Mohimani H, Morsy M, Moyne O, Neumann S, Neuweger H, Nguyen NH, Nothias-Esposito M, Paolini J, Phelan VV, Pluskal T, Quinn RA, Rogers S, Shrestha B, Tripathi A, van der Hooft JJJ, Vargas F, Weldon KC, Witting M, Yang H, Zhang Z, Zubeil F, Kohlbacher O, Böcker S, Alexandrov T, Bandeira N, Wang M, Dorrestein PC. 2020. Feature-based molecular networking in the GNPS analysis environment. Nat Methods 17:905–908.

56. Dettmer K, Aronov PA, Hammock BD. 2007. Mass spectrometry-based metabolomics. Mass Spectrom Rev 26:51–78.

57. Goulitquer S, Potin P, Tonon T. 2012. Mass spectrometry-based metabolomics to elucidate functions in marine organisms and ecosystems. Mar Drugs 10:849–880.

58. Gratton J, Phetcharaburanin J, Mullish BH, Williams HRT, Thursz M, Nicholson JK, Holmes E, Marchesi JR, Li JV. 2016. Optimized Sample Handling Strategy for Metabolic Profiling of Human Feces. Anal Chem 88:4661–4668.

59. Costea PI, Zeller G, Sunagawa S, Pelletier E, Alberti A, Levenez F, Tramontano M, Driessen M, Hercog R, Jung F-E, Kultima JR, Hayward MR, Coelho LP, Allen-Vercoe E, Bertrand L, Blaut M, Brown JRM, Carton T, Cools-Portier S, Daigneault M, Derrien M, Druesne A, de Vos WM, Finlay BB, Flint HJ, Guarner F, Hattori M, Heilig H, Luna RA, van Hylckama Vlieg J, Junick J, Klymiuk I, Langella P, Le Chatelier E, Mai V, Manichanh C, Martin JC, Mery C, Morita H, O’Toole PW, Orvain C, Patil KR, Penders J, Persson S, Pons N, Popova M, Salonen A, Saulnier D, Scott KP, Singh B, Slezak K, Veiga P, Versalovic J, Zhao L, Zoetendal EG, Ehrlich SD, Dore J, Bork P. 2017. Towards standards for human fecal sample processing in metagenomic studies. Nat Biotechnol 35:1069–1076.

60. Zhang X, Figeys D. 2019. Perspective and Guidelines for Metaproteomics in Microbiome Studies. J Proteome Res 18:2370–2380.

61. Turnbaugh PJ, Ley RE, Hamady M, Fraser-Liggett CM, Knight R, Gordon JI. 2007. The human microbiome project. Nature 449:804–810.

62. Integrative HMP (iHMP) Research Network Consortium. 2019. The Integrative Human Microbiome Project. Nature 569:641–648.

63. Nash AK, Auchtung TA, Wong MC, Smith DP, Gesell JR, Ross MC, Stewart CJ, Metcalf GA, Muzny DM, Gibbs RA, Ajami NJ, Petrosino JF. 2017. The gut mycobiome of the Human Microbiome Project healthy cohort. Microbiome 5:153.

64. Langille MGI, Zaneveld J, Caporaso JG, McDonald D, Knights D, Reyes JA, Clemente JC, Burkepile DE, Vega Thurber RL, Knight R, Beiko RG, Huttenhower C. 2013. Predictive functional profiling of microbial communities using 16S rRNA marker gene sequences. Nat Biotechnol 31:814–821.

65. Caporaso JG, Kuczynski J, Stombaugh J, Bittinger K, Bushman FD, Costello EK, Fierer N, Peña AG, Goodrich JK, Gordon JI, Huttley GA, Kelley ST, Knights D, Koenig JE, Ley RE, Lozupone CA, McDonald D, Muegge BD, Pirrung M, Reeder J, Sevinsky JR, Turnbaugh PJ, Walters WA, Widmann J, Yatsunenko T, Zaneveld J, Knight R. 2010. QIIME allows analysis of high-throughput community sequencing data. Nat Methods 7:335–336.

66. Integrative HMP (iHMP) Research Network Consortium. 2014. The Integrative Human Microbiome Project: dynamic analysis of microbiome-host omics profiles during periods of human health and disease. Cell Host Microbe 16:276–289.

67. Lloyd-Price J, Mahurkar A, Rahnavard G, Crabtree J, Orvis J, Hall AB, Brady A, Creasy HH, McCracken C, Giglio MG, McDonald D, Franzosa EA, Knight R, White O, Huttenhower C. 2017. Strains, functions and dynamics in the expanded Human Microbiome Project. Nature 550:61–66.

68. Human Microbiome Project Consortium. 2012. Structure, function and diversity of the healthy human microbiome. Nature 486:207–214.

69. earthmicrobiome. https://earthmicrobiome.org/. Retrieved 5 April 2024.

70. Thompson LR, Sanders JG, McDonald D, Amir A, Ladau J, Locey KJ, Prill RJ, Tripathi A, Gibbons SM, Ackermann G, Navas-Molina JA, Janssen S, Kopylova E, Vázquez-Baeza Y, González A, Morton JT, Mirarab S, Zech Xu Z, Jiang L, Haroon MF, Kanbar J, Zhu Q, Jin Song S, Kosciolek T, Bokulich NA, Lefler J, Brislawn CJ, Humphrey G, Owens SM, Hampton-Marcell J, Berg-Lyons D, McKenzie V, Fierer N, Fuhrman JA, Clauset A, Stevens RL, Shade A, Pollard KS, Goodwin KD, Jansson JK, Gilbert JA, Knight R, Earth Microbiome Project Consortium. 2017. A communal catalogue reveals Earth’s multiscale microbial diversity. Nature 551:457–463.

71. Gilbert JA, Jansson JK, Knight R. 2014. The Earth Microbiome project: successes and aspirations. BMC Biol 12:69.

72. Aksenov AA, da Silva R, Knight R, Lopes NP, Dorrestein PC. 2017. Global chemical analysis of biology by mass spectrometry. Nature Reviews Chemistry 1:1–20.

73. Fiehn O. 2016. Metabolomics by Gas Chromatography-Mass Spectrometry: Combined Targeted and Untargeted Profiling. Curr Protoc Mol Biol 114:30.4.1–30.4.32.

74. Wang Y, Xu L, Shen H, Wang J, Liu W, Zhu X, Wang R, Sun X, Liu L. 2015. Metabolomic analysis with GC-MS to reveal potential metabolites and biological pathways involved in Pb &Cd stress response of radish roots. Sci Rep 5:18296.

75. Kiseleva O, Kurbatov I, Ilgisonis E, Poverennaya E. 2021. Defining Blood Plasma and Serum Metabolome by GC-MS. Metabolites 12.

76. Alseekh S, Aharoni A, Brotman Y, Contrepois K, D’Auria J, Ewald J, C Ewald J, Fraser PD, Giavalisco P, Hall RD, Heinemann M, Link H, Luo J, Neumann S, Nielsen J, Perez de Souza L, Saito K, Sauer U, Schroeder FC, Schuster S, Siuzdak G, Skirycz A, Sumner LW, Snyder MP, Tang H, Tohge T, Wang Y, Wen W, Wu S, Xu G, Zamboni N, Fernie AR. 2021. Mass spectrometry-based metabolomics: a guide for annotation, quantification and best reporting practices. Nat Methods 18:747–756.

77. Plumb RS, Gethings LA, Rainville PD, Isaac G, Trengove R, King AM, Wilson ID. 2023. Advances in high throughput LC/MS based metabolomics: A review. Trends Analyt Chem 160:116954.

78. Schönberger K, Mitterer M, Glaser K, Stecher M, Hobitz S, Schain-Zota D, Schuldes K, Lämmermann T, Rambold AS, Cabezas-Wallscheid N, Buescher JM. 2023. LC-MS-Based Targeted Metabolomics for FACS-Purified Rare Cells. Anal Chem 95:4325–4334.

79. Xiao JF, Zhou B, Ressom HW. 2012. Metabolite identification and quantitation in LC-MS/MS-based metabolomics. Trends Analyt Chem 32:1–14.

80. Zhou B, Xiao JF, Tuli L, Ressom HW. 2012. LC-MS-based metabolomics. Mol Biosyst 8:470–481.

81. Peng Y, Zhang Z, He L, Li C, Liu M. 2024. NMR spectroscopy for metabolomics in the living system: recent progress and future challenges. Anal Bioanal Chem 416:2319–2334.

82. Emwas A-H, Roy R, McKay RT, Tenori L, Saccenti E, Gowda GAN, Raftery D, Alahmari F, Jaremko L, Jaremko M, Wishart DS. 2019. NMR Spectroscopy for Metabolomics Research. Metabolites 9.

83. Moco S. 2022. Studying Metabolism by NMR-Based Metabolomics. Front Mol Biosci 9:882487.

84. Putri SP, Fukusaki E. 2016. Mass Spectrometry-Based Metabolomics: A Practical Guide. CRC Press.

85. Zhang A, Wang W. 2021. Mass Spectrometry-Based Metabolomics in Clinical and Herbal Medicines: Strategies, Technologies, and Applications. John Wiley & Sons.

86. Raftery D. 2014. Mass Spectrometry in Metabolomics: Methods and Protocols. Humana.

87. Ramautar R. 2021. Advanced Mass Spectrometry-based Analytical Separation Techniques for Probing the Polar Metabolome. Royal Society of Chemistry.

88. González-Domínguez R. 2022. Mass Spectrometry for Metabolomics. Springer Nature.

89. Peters K, Bradbury J, Bergmann S, Capuccini M, Cascante M, de Atauri P, Ebbels TMD, Foguet C, Glen R, Gonzalez-Beltran A, Günther UL, Handakas E, Hankemeier T, Haug K, Herman S, Holub P, Izzo M, Jacob D, Johnson D, Jourdan F, Kale N, Karaman I, Khalili B, Emami Khonsari P, Kultima K, Lampa S, Larsson A, Ludwig C, Moreno P, Neumann S, Novella JA, O’Donovan C, Pearce JTM, Peluso A, Piras ME, Pireddu L, Reed MAC, Rocca-Serra P, Roger P, Rosato A, Rueedi R, Ruttkies C, Sadawi N, Salek RM, Sansone S-A, Selivanov V, Spjuth O, Schober D, Thévenot EA, Tomasoni M, van Rijswijk M, van Vliet M, Viant MR, Weber RJM, Zanetti G, Steinbeck C. 2019. PhenoMeNal: processing and analysis of metabolomics data in the cloud. Gigascience 8.

90. Schmid R, Heuckeroth S, Korf A, Smirnov A, Myers O, Dyrlund TS, Bushuiev R, Murray KJ, Hoffmann N, Lu M, Sarvepalli A, Zhang Z, Fleischauer M, Dührkop K, Wesner M, Hoogstra SJ, Rudt E, Mokshyna O, Brungs C, Ponomarov K, Mutabdžija L, Damiani T, Pudney CJ, Earll M, Helmer PO, Fallon TR, Schulze T, Rivas-Ubach A, Bilbao A, Richter H, Nothias L-F, Wang M, Orešič M, Weng J-K, Böcker S, Jeibmann A, Hayen H, Karst U, Dorrestein PC, Petras D, Du X, Pluskal T. 2023. Integrative analysis of multimodal mass spectrometry data in MZmine 3. Nat Biotechnol 41:447–449.

