## Supplementary Figures for "S-Wipe: stool sample collection for metabolomic gut health tracking"

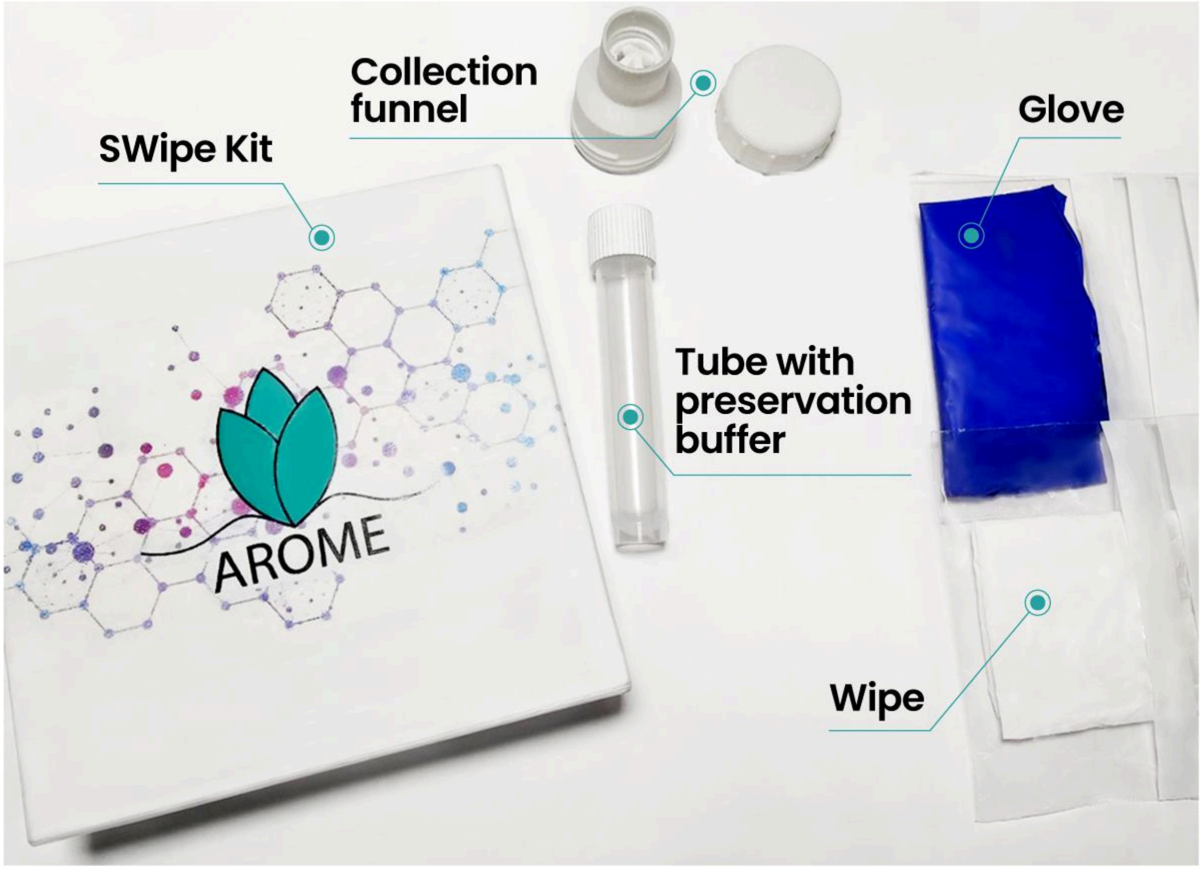


Figure 1. S-Wipe kit.

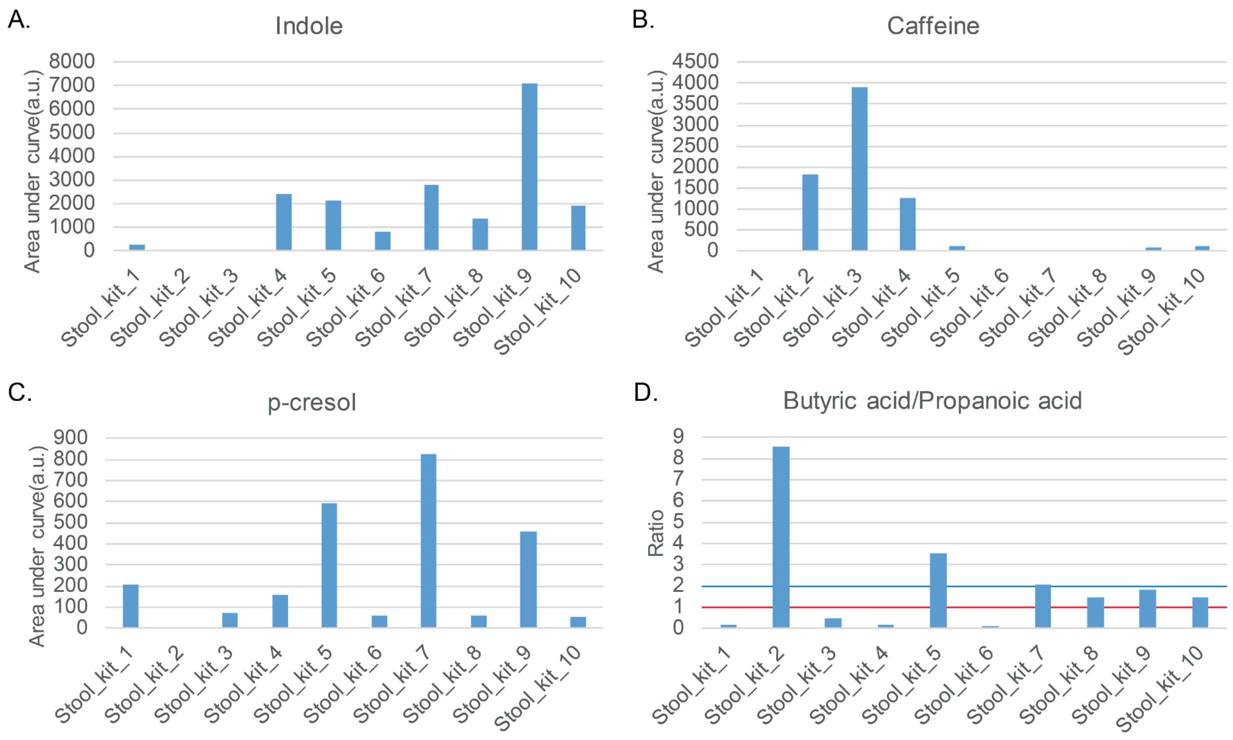


Figure 2. A. Indole, major stool smell component B. Caffeine, only found in coffee/tea consumers. C. p-cresol, a breakdown product of protein and proposed biomarker of autism in kids (67–72). D. Excessively low ratio of butyrate to propionate (subjects below red bar) is associated with insufficient fiber intake, loss of butyrate producing bacteria, inflammatory conditions (42, 43)(43–46), while excessively high ratio (subjects above blue bar) may be indicative of SIBO (43–45). The expected ratio for a healthy population is 1-2 (42, 43, 45–48).

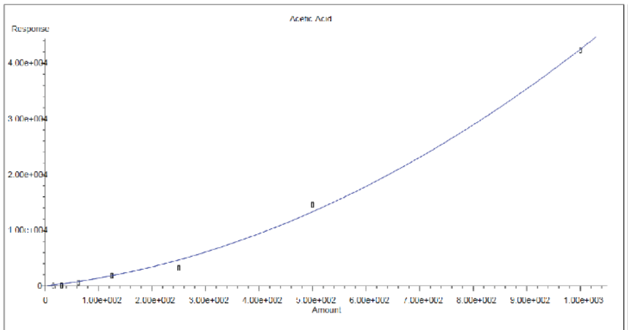


R = 3.17e-002 A*A + 1.08e+001 A + 0.00e+000
Coef of Det (r^2) = 0.998 Curve Fit: Quad/(0,0)

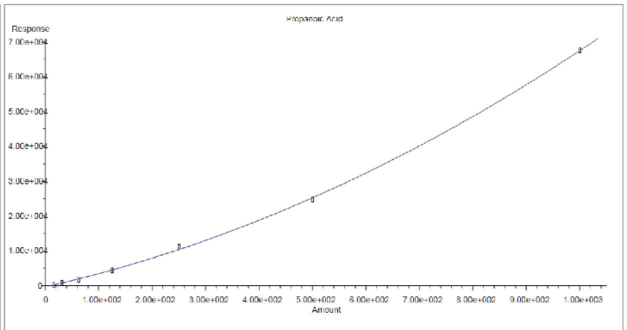


R = 3.34e-002 A*A + 3.44e+001 A - 3.05e+002
Coef of Det (r^2) = 1.000 Curve Fit: Quadratic

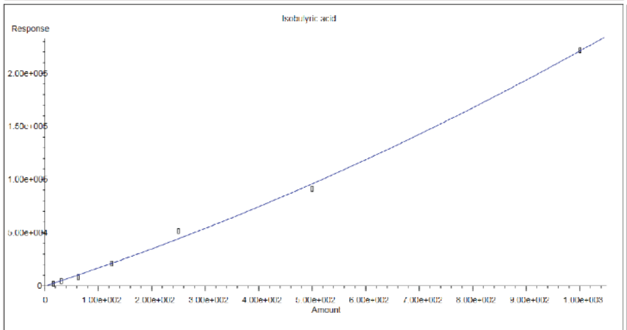


R = 5.75e-002 A*A + 1.64e+002 A - 2.43e+002
Coef of Det (r^2) = 0.998 Curve Fit: Quadratic

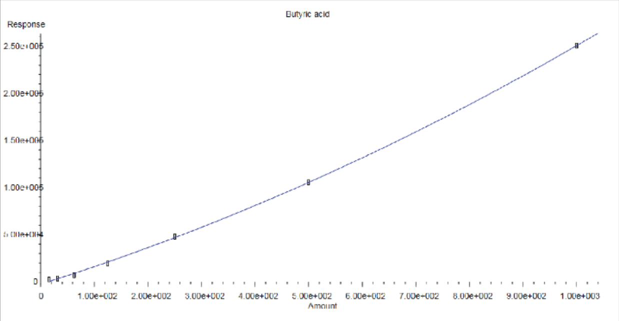


R = 7.42e-002 A*A + 1.79e+002 A - 2.58e+003
Coef of Det (r^2) = 1.000 Curve Fit: Quadratic

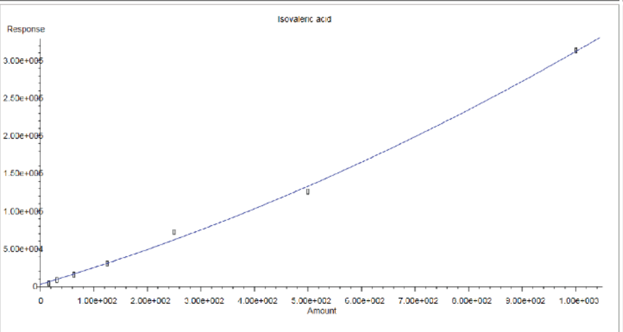


R = 9.72e-002 A*A + 2.12e+002 A + 2.91e+003
Coef of Det (r^2) = 0.998 Curve Fit: Quadratic

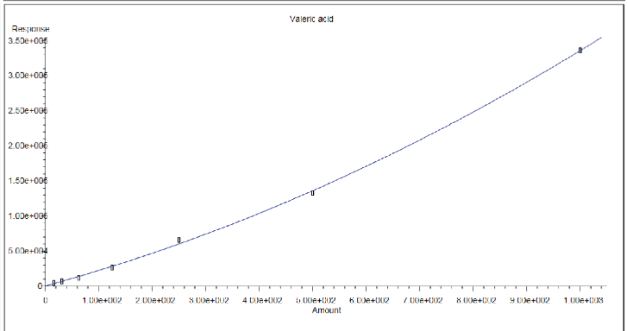


R = 1.28e-001 A*A + 2.08e+002 A + 2.52e+002
Coef of Det (r^2) = 0.999 Curve Fit: Quadratic

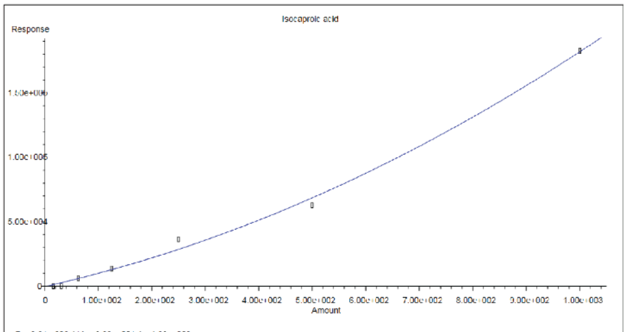


R = 8.91e-002 A*A + 9.29e+001 A - 1.68e+002
Coef of Det (r^2) = 0.996 Curve Fit: Quadratic

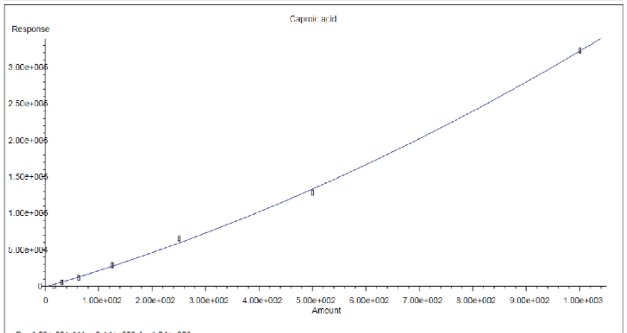


R = 1.09e-001 A*A + 2.14e+002 A - 1.04e+003
Coef of Det (r^2) = 0.999 Curve Fit: Quadratic

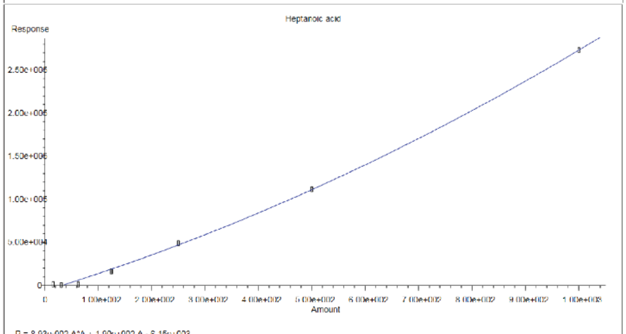


R = 8.93e-002 A*A + 1.90e+002 A - 6.15e+003
Coef of Det (r^2) = 0.999 Curve Fit: Quadratic

Figure 3. Calibration curves for short and medium chain fatty acids. Physiological levels are extensively covered by quantitative linearity for these molecules.

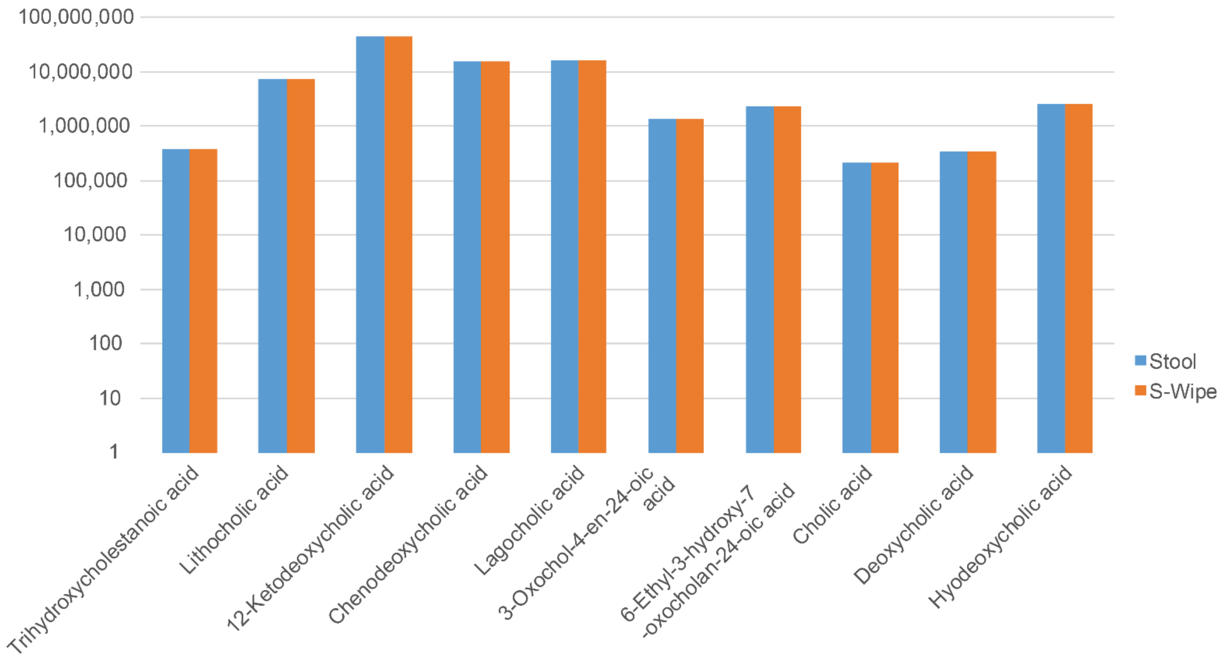


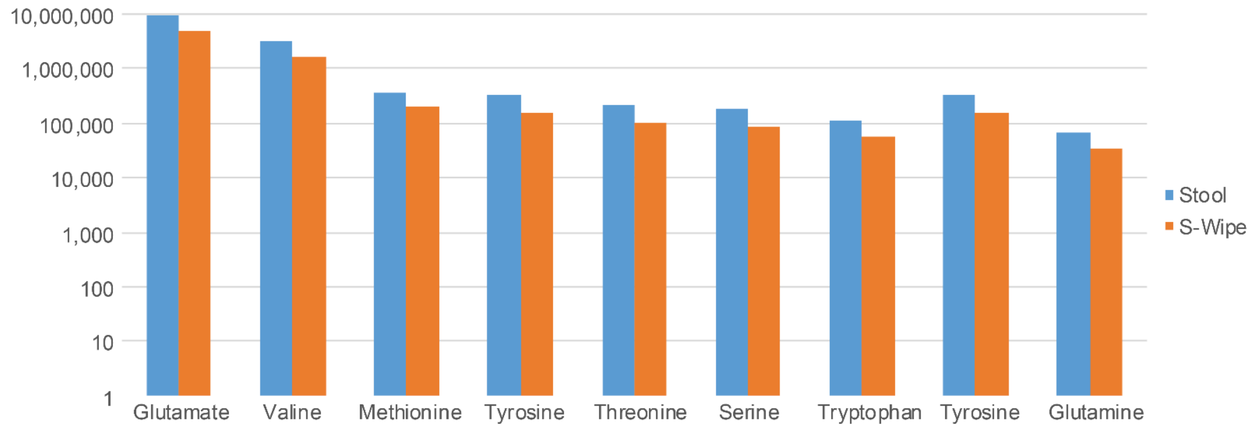


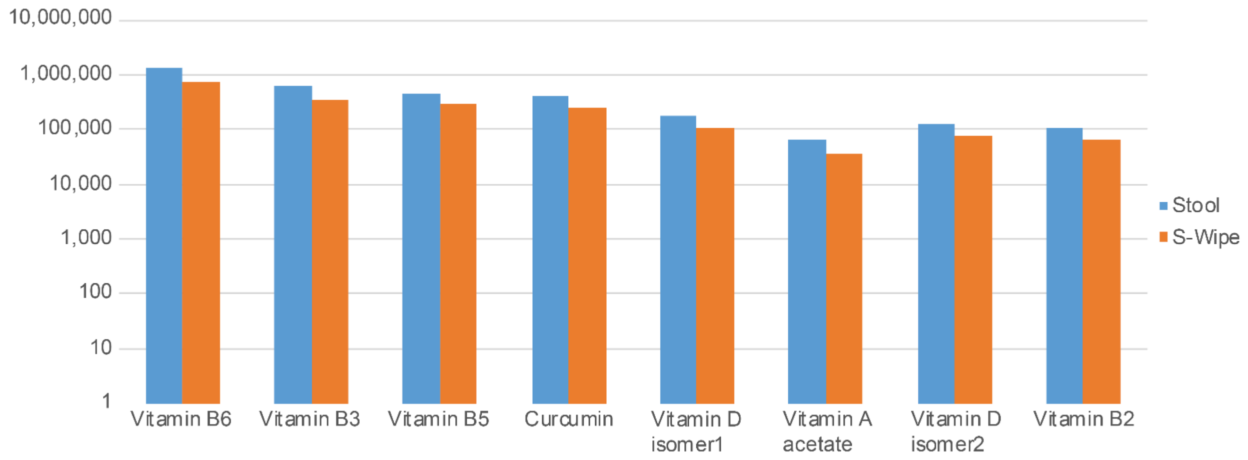

Figure 4. Examples of amounts of various metabolites detected with a variety of protocols(see Protocols study in methods). Bile acids, amino acids and vitamins. The abundance of the captured metabolites for both regular direct stool collection and S-Wipe methods are similar.

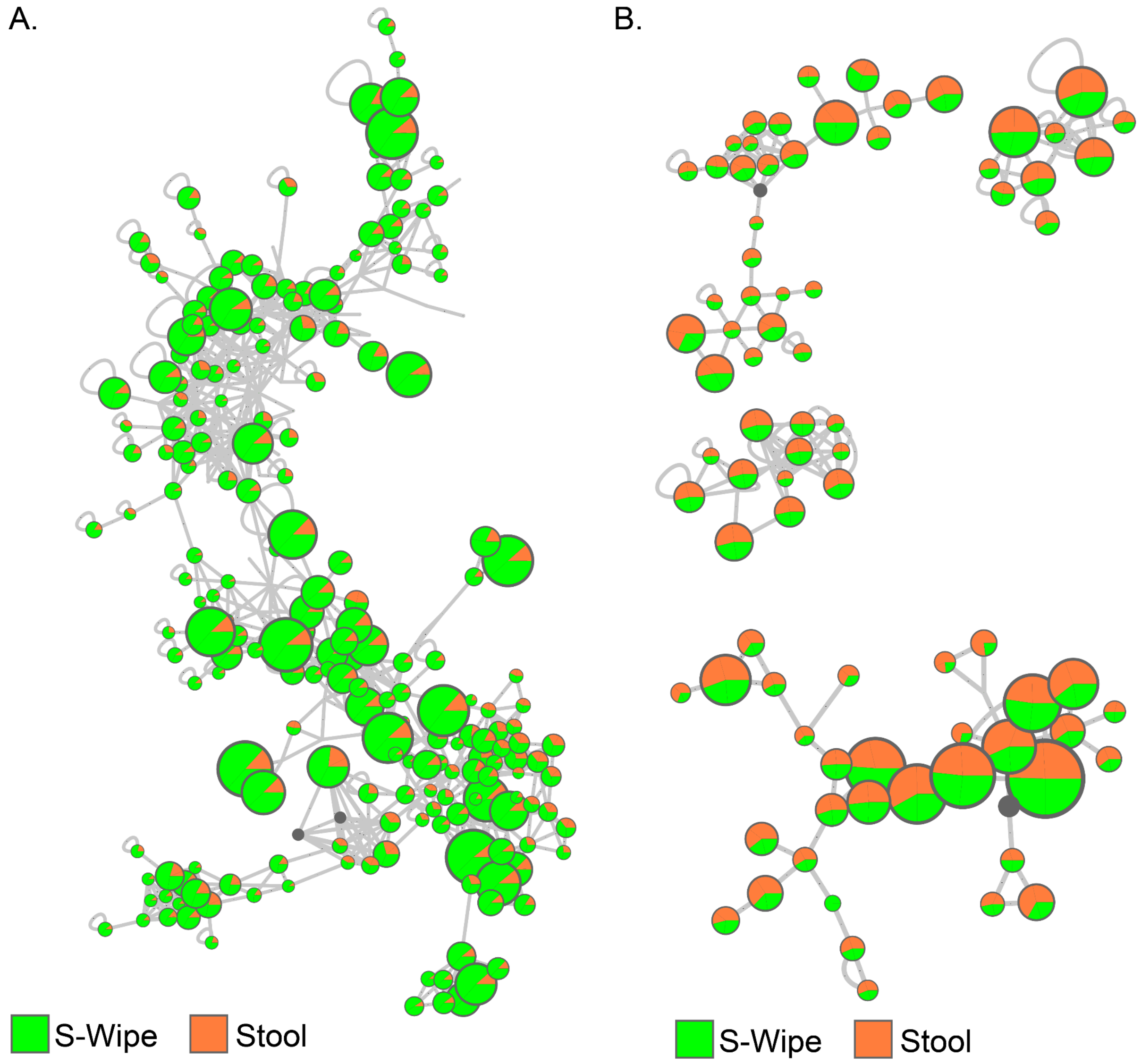


Figure 5. LC-MS/MS network clusters of S-wipe vs regularly extracted stool samples. A) Cluster of polyethylene glycol contaminants from wipes. B) Clusters identified as those of stool endogenous molecules. Each node represents the presence of specific metabolites.
