## Supplementary material for "S-Wipe: stool sample collection for metabolomic gut health tracking": Normalized Density

| | Normalized Density | Video Snippet |
|---|---|---|
| **Sample-1** | 376.1389528 | V1 |
| **Sample-2** | 595.9718628 | V2 |
| **Sample-3** | 544.7059816 | V3 |
| **Sample-10** | 86.54310438 | V10 |
| **Sample-11** | 214.1285668 | V11 |
| **Sample-12** | 111.2916387 | V12 |
